# Late flowering and low synchrony with males enhance female reproductive success in the dioecious palm *Chamaedorea pinnatifrons*

**DOI:** 10.1101/057547

**Authors:** Luis D. Ríos, G. Barrantes, Alfredo Cascante-Marin

## INTRODUCTION

Dioecious plants (i.e., with separate sexes) obligatorily require the contribution of a conspecific, and the synchronization of a series of events to achieve sexual reproduction (Bawa 1980). In these plants pollination is only possible when a series of successive events occur: temporal overlap of male and female flowering, sufficient quality pollen is produced by male plants, there is an effective mechanism for pollen capture in female plants, and adequate pollen transportation (Givnish 1982). However, pollination could be farther affected by population density, population sex ratio, individual phenological behavior, individual size, and the species pollination system (Larson and Barrett 2000; Knight et al. 2005). Usually, plants show a low reproductive success (measured as fruit production or fruit set) at decreasing population densities (Knight et al. 2005). A low population density may also increase the distance between male and female plants, reducing pollen availability for the last, and in consequence hampering their reproductive success (Steven and Waller 2006).

The sex ratio in populations of dioecious plants often deviates from a 1:1 sex ratio, presenting either an excess of males or females (Field et al. 2013). Skewed sex ratios might reduce the fitness of individuals in the population (Fisher 1930). For example, Shelton (2008) showed that seed production in two dioecious sea grass species was lower in populations that had a sex ratio skewed towards females. On the contrary, Carlsson-Graner et al. (1998) found that females of *Silene dioica* (Caryophyllaceae) in male biased populations experienced an increased rate of pollen deposition per female flower. It is then expected that with decreasing male abundance, female reproductive success will become increasingly pollen limited (House 1992). Similarly, it could be expected that in populations with an excess of females, competition between each other for pollen availability should play a role in their reproductive success.

Variation in the reproductive phenology can also affect an individual's reproductive success (Elzinga et al. 2007). Munguía-Rosas et al. (2011) through a meta-analysis showed that selection usually favors early flowering individuals. However, in plants with strict outbreeding reproduction (e.g., dioecious species) a high degree of flowering synchrony is expected for effective cross-pollination between the sexes (Pires et al. 2013). Early or late flowering plants could experience a functional low population density, and/or a highly skewed sex ratio, which in return will affect negatively their fruit production. Therefore, it is expected that selection will favor female plants that are able to flower when mate availability is higher (e.g., Augspurger 1983), and that plant reproductive success is affected by the interaction among phenology, population density and sex ratio.

Another factor that may affect fruit production is the plant's pollination system (Fenster et al. 2004). Plants vary in their degree of pollination specialization, from those specialist plants, pollinated by only one or few pollinators, to generalist species, pollinated by many different taxa (Faegri and van der Pijl 1979). Within a community of flowering plants, most species are usually generalists, so that some degree of competition for the same pool of pollinators is expectable (Petanidou et al. 2014). For instance, Runquist (2013) found that the generalist species *Limnanthes douglasii* (Limnanthaceae) experiences significant pollen limitation during the early stages of the flowering season, and this limitation increases with the number of co-flowering species in the community. Specialist plants could also be limited by pollinators, but with a weaker or null effect of co-flowering plants.

In this paper we analyze the effect of phenology, male and female neighborhood, size, and floral display on the reproductive success of the understory dioecious palm *Chamaedorea pinnatifrons* (Arecaceae) in a Costa Rican montane forest. This palm is pollinated by wind and a single species of thrips (Thysanoptera: Thripidae) in a mixed pollination system (Rios et al. 2014). Thrips use male inflorescences for feeding and reproduction, and pollinate female flowers by deception, most likely by the similarity in fragrance composition. The pollination system of *C. pinnatifrons* seems to be highly specialized, as the thrips species has been recorded only in flowers of *Chamaedorea* (Porter Morgan 2007; Rios et al. 2014). Based on this, we also related female reproductive success to the availability of insects throughout the flowering season. Specifically, our objectives in this study were: 1) to describe the phenology of a population of *C. pinnatifrons* in a Costa Rican montane forest, 2) to determine the effect of phenology, size, density and distance to the nearest conspecific on the female reproductive success (measured as fruit production or fruit set) and, 3) to relate our findings on female reproductive success with the abundance of pollinators.

## MATERIALS AND METHODS

### Study species and site

*Chamaedorea pinnatifrons* (subgenus *Chamaedorea)* is an understory palm and occurs between southern Mexico to Bolivia, from 400 to 2600 m a.s.l. (Hodel 1992). Individuals have three to eight pinnate leaves and develop a single stem up to 4 m high (Hodel 1992, Grayum 2003). Seeds are dispersed by birds (Orozco-Segovia et al. 2003) and a curculionid beetle (similar to the one found by Oyama (1991) at Los Tuxtlas, Mexico; Rios, unpbl. data) predate pre-dispersal seeds. Ataroff and Schwarzkopf (1994) estimated that males could live up to 61 years, while females up to 66 years at a Venezuelan cloud forest.

Field work was conducted in La Carpintera montane cloud forest in the Costa Rican Central Valley, (9°53’20” N; 83°58’10” W). The site consists of an elevated mountainous terrain rising from 1500 to 1800 m a.s.l., and includes a forest fragment of nearly 2400 ha that covers the ridge and highest portion of the mountain slopes. The forest consists of mainly old secondary forest (>50 years old) interspersed with older remnant forest patches, that include large trees such as oaks *(Quercus* spp.), fig trees *(Ficus* spp.), and several species of Lauraceae (Sánchez et al. 2008). Mean annual precipitation is 1839 mm and mean annual temperature is 16.1 °C. The annual rainfall follows a seasonal pattern, with a period of low precipitation ( < 60 mm per month) from December to April (IMN, undated).

### Experimental design

#### Study plot, population phenology and female reproductive success

We established a 40 × 40 m plot at La Carpintera. Within the study plot, we labeled all reproductive individuals (88 males and 94 females) with aluminum labels, and for each individual we counted the number of leaves, measured their height (only females), and mapped their position in a X-Y coordinate system (Fig. S1). We registered the flowering phenology of each individual every 3-4 days from February to July 2012, for a total of 41 visits. We used this censuses period because previous observations showed that anthesis of inflorescences lasted 3-7 days. During each census, we assigned each inflorescence to one of the following phenophases: (1) inflorescences enclosed by the peduncular bracts, (2) inflorescences emerged from the bracts and flowers in bud, (3) flowers in anthesis (male: releasing pollen, female: stigma receptive, green and bright), (4) flowers in post-anthesis (male: flowers senescent, female: stigma dry and brown) and, (5) fruits green (inmature) or black (mature). Female reproductive success was estimated as the number of fruits produced per inflorescence, and by the fruit set (fruits/flowers) per inflorescence of 115 inflorescences (n = 74 individuals).

#### Data analysis

We evaluated the effect of variables that correlate with plant vigor (i.e., stem height, number of leaves), male neighborhood (i.e., nearest synchronous male, number of synchronous males at five and ten meters around the female plant), female neighborhood (i.e., nearest synchronous female, number of synchronous females at five and ten meters around the focal female plant), number of flowers in each inflorescence, flowering synchrony (Marquis 1988) within and among sexes (equation in Supporting Information), flowering date, and the interaction between synchrony and flowering date, on the number of fruits and fruit set per inflorescence. We used General Linear Mixed Models (GLMM) with the predictor variables and the inflorescence nested within the individuals as random factors. For each response variable we selected a set of possible models, including a model without random effects, and then chose the optimal model using the Akaike information criteria (AIC;Zuur et al. 2009). All analyses were performed with the R statistical language, version 3.2.3 (R Core Team 2015).

We also analyzed the frequency of fruit production throughout the flowering season by dividing the season by terciles. Each tercile represented different stages in the season: early flowering (n = 34 female inflorescences), flowering peak (n = 39) and, late flowering (n = 42). We later plotted the fruit production frequency for each tercile.

### Thrips population

We estimated thrips abundance by sampling insects directly from staminate inflorescences in anthesis. From early March to mid-May (11 weeks), insects were sampled from three staminate inflorescences per week (*n* = 33 inflorescences) haphazardly chosen outside the study plot. Insects were collected by placing the inflorescences in a plastic bag and gently shaking it to detach insects, which were later preserved in 70% ethanol. We counted adult thrips in the laboratory with a stereoscope. The amount of thrips per inflorescence was divided by the number of rachillae to correct for the size of the inflorescence, as the amount of thrips caught might be influenced by the size of the sampling unit (Lewis 1997). Data is presented as the mean (± standard error) number of thrips/rachillae for each week. Voucher specimens were deposited in the Museum of Zoology, Universidad de Costa Rica.

## RESULTS

### Phenology and reproductive success

Flowering of *C. pinnatifrons* lasted five months: it began by early February, had a peak by late march and early April (prior to the rainy season), and ended by early July (Fig.1). Mean fruit production was 99.7 (± 12.1 SE, min = 0, max = 735). The number of fruits per inflorescence was best predicted by the model that included all predictor variables with inflorescence nested within individual as random factor. The amount of fruits produced was significantly higher in inflorescences from individuals with shorter stems, that had more flowers per inflorescence, and which had lower flowering synchrony with males and flowered late in the season (Table 1; Fig. 2).

**Fig. 1.**
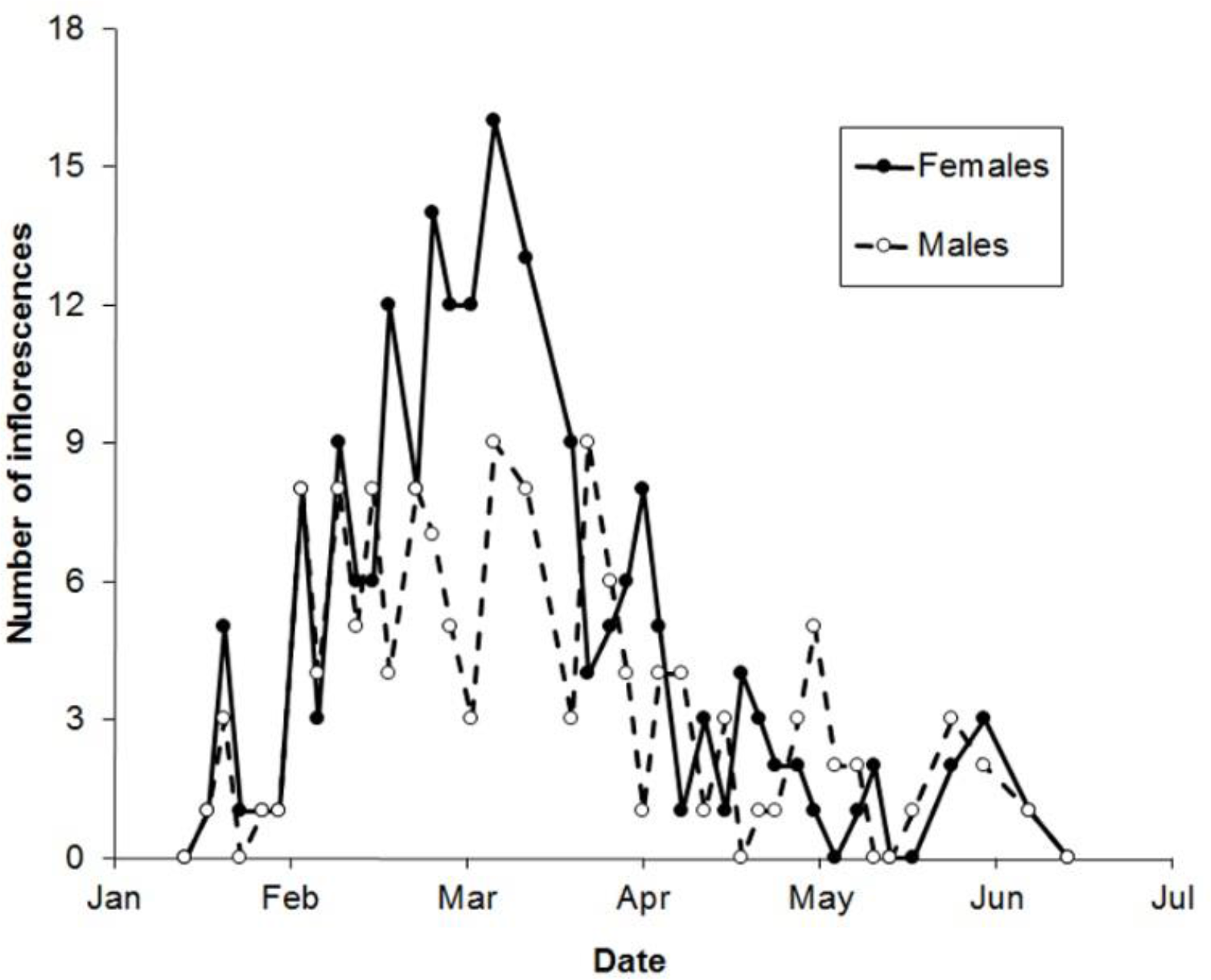
Reproductive phenology of *Chamaedorea pinnatifrons* at La Carpintera, Costa Rica, during the 2012 season. Filled circles indicate female inflorescences in anthesis and open circles indicate male inflorescences in anthesis.

**Table 1.** Effect of predictor variables (see table content) on number of fruits and fruit set (fruits/flowers) based on General Linear Mixed Models. Number of fruits was best predicted by a model that included six predictor variables with inflorescence nested within individual plant as random factors. Fruit set was best predicted by a model that included five variables with inflorescence nested within individual plant as random factors. Variables with a significant effect are highlighted in bold.

| Response variable: Number of fruits |  |  |  |  |
| --- | --- | --- | --- | --- |
| Effect | Coefficient | SE | T | P |
| Intercept | -47.37 | 81.05 | -0.58 | 0.5608 |
| **Stem height** | **-0.48** | **0.17** | **-2.87** | **0.0054** |
| No. leaves | 22.87 | 11.95 | 1.91 | 0.0597 |
| **No. flowers** | **0.18** | **0.04** | **5.05** | **<0.00001** |
| Nearest Male | -1.80 | 1.08 | -1.66 | 0.1075 |
| Males at 5 m | -25.62 | 16.00 | -1.60 | 0.1197 |
| Males at 10 m | 3.57 | 8.31 | 0.43 | 0.6704 |
| Nearest female | -1.18 | 1.18 | -1.00 | 0.3260 |
| Females at 5 m | 1.98 | 11.49 | 0.17 | 0.8640 |
| Females at 10 m | -13.68 | 11.03 | -1.24 | 0.2245 |
| **Synchrony males** | **-17220.13** | **6944.06** | **-2.48** | **0.0190** |
| **Flowering date** | **6.51** | **1.57** | **4.13** | **0.0003** |
| **Male sync\*date** | **1156.34** | **501.44** | **2.31** | **0.0282** |
| Response variable: Fruit set |  |  |  |  |
| Effect | Coefficient | SE | T | P |
| Intercept | 0.16 | 0.10 | 1.65 | 0.1030 |
| **Stem height** | **-0.00** | **0.00** | **-3.53** | **0.0007** |
| No. leaves | 0.03 | 0.00 | 1.81 | 0.0749 |
| Nearest Male | -0.00 | 0.00 | -1.69 | 0.1014 |
| Males at 5 m | -0.03 | 0.02 | -1.69 | 0.1003 |
| Males at 10 m | 0.01 | 0.01 | 0.73 | 0.4698 |
| Nearest Female | -0.00 | 0.00 | -1.31 | 0.1975 |
| Females at 10 m | -0.02 | 0.01 | -1.40 | 0.1699 |
| **Synchrony males** | **25.52** | **8.70** | **-2.93** | **0.0062** |
| **Flowering date** | **0.01** | **0.00** | **4.82** | **<0.00001** |
| **Male sync\*date** | **1.44** | **0.63** | **2.29** | **0.0286** |

**Fig. 2.**
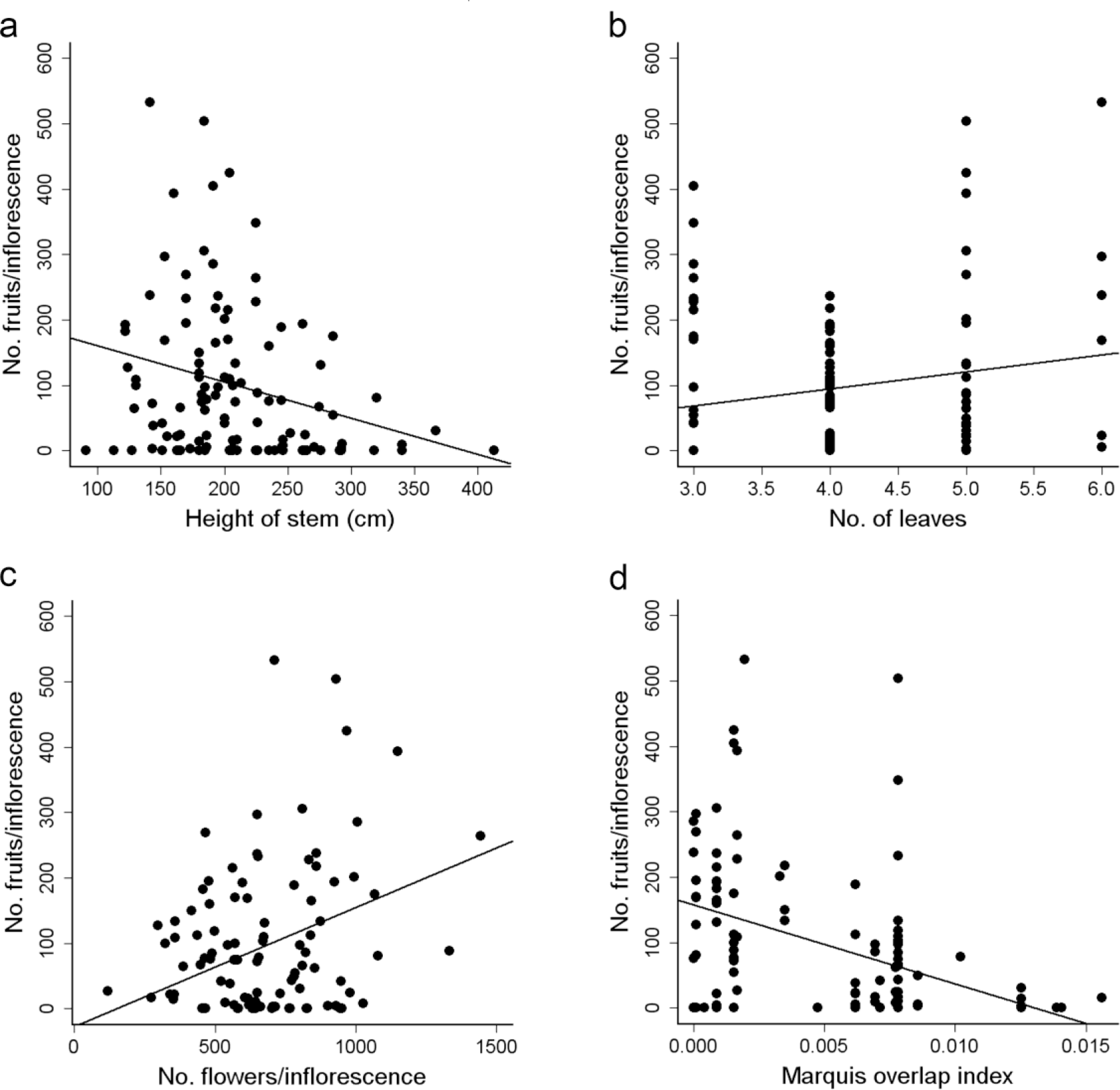
Variables that have a significant effect on number of fruits. Number of fruits/inflorescence decreases with height of the stem (a), increases with number of leaves (b), and number of flowers on the inflorescence (c), and decreases with the overlap with male flowering-plants (c).

Mean fruit set was 0.144 (± 0.016, min = 0, max = 0.832). Fruit set was also significantly higher in inflorescences from lower stems, with lower male synchrony and which had a late flowering date (Table 1). In this case, fruit set was best predicted by a model that included the same variables, but females within 5 m from the focal female plant, with inflorescence nested within individual as random factor. For both response variables, number of fruits and fruit set, flowering date and the synchrony with male flowering plants had a significant interaction (Table 1, Fig. S2). Lower values of synchrony occurred toward the end of the flowering season (Fig. S2). Synchrony with males and flowering date correlated positively with number of fruits (r = 0.59, p < 0.00001) and fruit set (r = 0.63, p < 0.001), indicating that plants produced more fruits and had higher fruit sets at the end of the flowering season.

In general, 35% (40/115) of all the inflorescences set less than 10 fruits. Of these inflorescences, 29 didn't produce any fruit. Thirty percent (35/115) produced between 11 and 100 fruits, while the remaining 35% of the inflorescences produced more than 100 fruits each. When the inflorescences were categorized by their flowering phenology, most of the early flowering inflorescences (65%, 22/34) produced less than 10 fruits each, and only 6% (2/34) set more than 100 (Fig. 3). In contrast, 59% (23/39) of the inflorescences that flowered during the population flowering peak produced more than 10 fruits. In a similar fashion, 74% (31/42) of the late flowering inflorescences set more than a 100 fruits each; meanwhile, only 7% (3/42) inflorescences produced less than 10 fruits (Fig. 3).

**Fig. 3.**
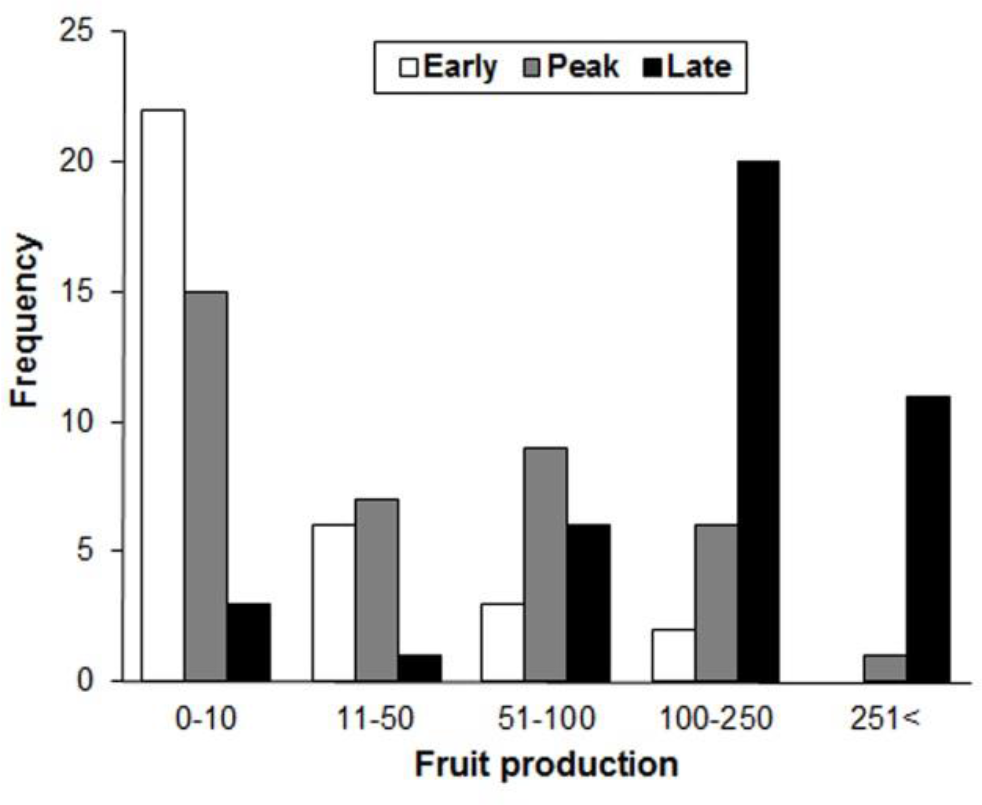
Fruit production of 115 female inflorescences of *Chamaedorea pinnatifrons* at La Carpintera Cloud Forest, ordered according to the inflorescence’s flowering phenology: early flowering, flowering peak or late flowering

### Thrips population

The number of adult thrips in staminate inflorescences in anthesis increased steadily as the flowering season progressed (Fig. 4). We found a mean of 1044 ± 145 (±SE, min = 35, max = 2893) thrips/inflorescence. The mean amount of thrips sampled in each inflorescence increased 13-fold between early March and mid-May. In the first week of March 20.8 ± 10.6 thrips/rachillae were sampled from staminate inflorescences, whilst by the third week of May 282.0 ± 74.4 thrips/rachillae were collected (Fig. 4).

**Fig. 4.**
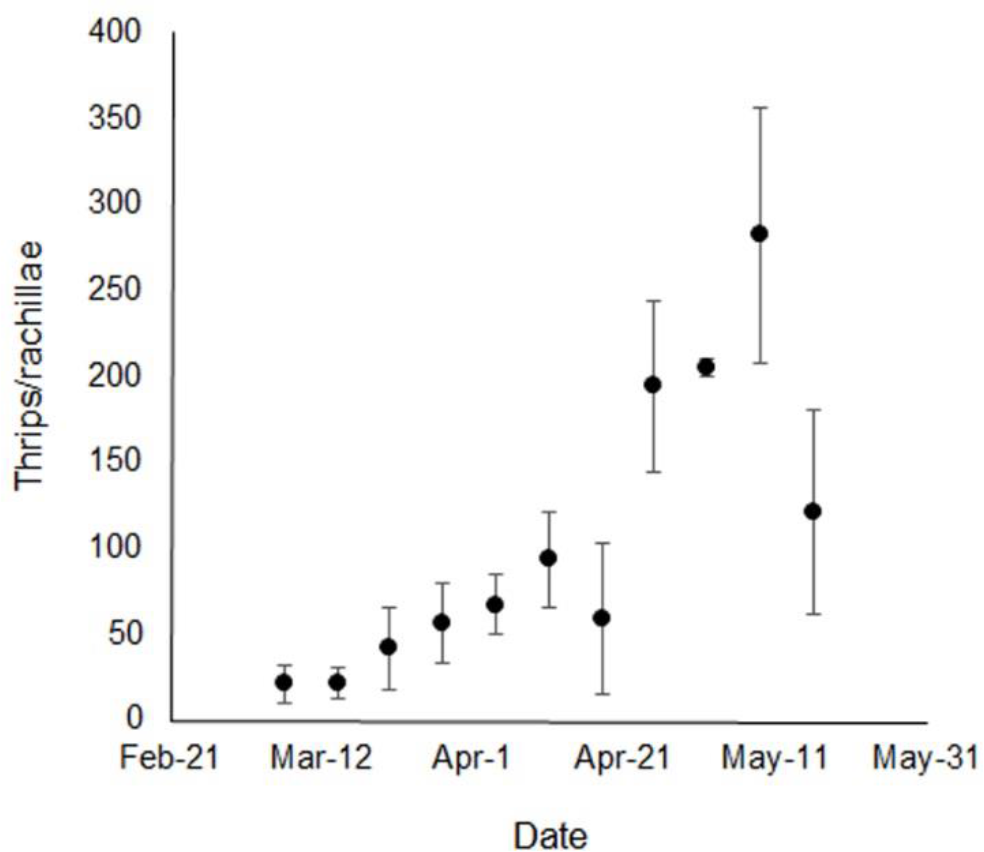
Adult thrips caught from staminate inflorescences of *C. pinnatifrons* (*n* = inflorescences) at La Carpintera, Costa Rica during the 2012 flowering season for 11 dates. Data is presented as the mean ± SE (vertical lines) thrips/rachillae of three inflorescences for each census date.

## DISCUSSION

Our findings show that the female reproductive success of *C. pinnatifrons* at La Carpintera was higher in late flowering females with low flowering synchrony with males, and which also had a high number of flowers per inflorescence and shorter stem height. Most studies that analyze the effect of phenology on fruit production have found that early flowering plants should be positively selected as they might experience less competition for pollinators or have more time for seed maturation (Munguía-Rosas et al. 2011). It is noteworthy that fruit output in *C. pinnatifrons* shows an opposite trend, in as much late flowering females with lower synchrony with males are the ones who presented the highest reproductive success. In addition, our results suggest that the reproductive success of *C. pinnatifrons* is potentially pollen limited as 25% (29/115) of all inflorescences didn't produce any fruit.

Many studies have found that presenting flowers during the population flowering peak should be positively selected as the amount of potential mates is higher at that particular time than in off-peak flowering (Elzinga et al. 2007). However, late flowering can be favored if pollinator reliability is low early in the season, and increases by the end of it. If the pollinator is highly specific and its population grows as the flowering season progresses, then early flowering plants would receive fewer visits by pollinators in comparison to late flowering plants. This translates to a higher reproductive success of late flowering individuals. A similar hypothesis has been invoked to explain the high visitation rate and later higher fruit onset that experience some species during the population flowering peak (Augspurger 1983). As demonstrated, thrips population increased along with the flowering season. Therefore, the higher fruit production of late flowering plants, might be partially explained by a potential increase in the insects' visitation rate to pistillate inflorescences. Although the data obtained in this manner was from staminate inflorescences, its reliability stems from the fact that thrips have been recorded visiting both kinds of inflorescences simultaneously at the study site (Rios et al. 2014).

Although late flowering females effectively captured more pollen than early or peak flowering plants, it is possible that the genetic diversity of the pollen they caught was lower, as less male plants were flowering during that period (exemplified in the lower male synchrony indexes). It has been suggested that multiple paternity within a fruit or infructescence could potentially increase the progeny's fitness in the form of seed mass or germination success (Pélabon et al. 2015). Thus far, the evidence gathered regarding such an expectation has been inconclusive (Pélabon et al. 2015) and it is likely that other selective forces might be shaping the population's phenology. To test this hypothesis, it would be necessary to compare the progeny's fitness (seedling emerge and/or dry weight) and the genetic diversity of the seeds produced during the flowering peak against the ones produced by early and late flowering inflorescences. This could potentially explain how such a distinctive population flowering peak would endure, even with an apparent lower reproductive success.

Another possibility is that the population's flowering phenology might be affected by pre-dispersal seed predation (Kolb et al. 2007; Archibald et al. 2012). Pre-dispersal seed predation on *C. pinnatifrons* at the study site is mediated by an unidentified curculionid beetle (Ríos Pers. Obs.). Brody and Mitchell (1997) showed that an increase in a plants' fruit output could enhance the rate of pre-dispersal seed predation. Although we were not able to quantify seed predation in this study, it is likely that the beetles are predating late flowering inflorescences more heavily, in as such they could represent a more attractive resource because of their higher fruit production. An integrated experiment in which the effect of both antagonistic (beetles) and mutualistic (thrips) selective agents are taken into account (e.g., Ehrlén 2014), could help elucidate the strength of the selective agents over the individuals' flowering phenology and test whether or not the populations' phenology will shift to a later mean flowering date.

The result that the number of flowers per inflorescence is positively correlated with fruit production is not surprising, as many studies have already described such a trend (Wyatt 1982; Thomson 1988; Oharam and Higashi 1994). Inflorescences with more flowers can be positively selected if they are capable of attracting a higher amount of pollinators than smaller ones (Wyatt 1982). Similarly, as wind also participates in the pollination of *C. pinnatifrons* (Rios et al. 2014), it can be expected that inflorescences with numerous flowers are capable of sampling a larger volume of the airstream, which in return maximizes pollen capture and its' reproductive success (Friedman and Barrett 2011).

The expectation that females with a larger size should have had a higher reproductive success was not met, as individuals with lower stem height had a significantly higher fruit production. We noted that height was negatively correlated with the number of leaves a palm had (R^2^ = −0.22, p < 0.05); thus it can be expected that the lower number of leaves that taller palms sustained might have negatively affected their reproductive capacity. In relation to this topic, Martínez-Ramos et al. (2009) found that females of *Chamaedorea elegans* with a greater leaf area had an improved fecundity. Less leaf area would render a low photosynthetic capacity, and in consequence affect the amount of available resources for reproduction (Svenning 2001). Also, it is likely that size and age might be positively correlated in *C. pinnatifrons*, therefore the decrease in the reproductive success might reflect a progressive loss of reproductive capacity through the individual's lifetime. In this regard, dendrochronological studies have found a negative association between trunk size and fruit production (Thomas 2011).

## Conclusion

Our experimental design shows that fruit production and fruit set (as proxy for reproductive success) is highly affected by the individual's flowering phenology. The result that late flowering individuals presented a greater fecundity is most likely an effect of the increase in pollinators abundance that occurs as the flowering seasons advance. This results contrast with previous research (Munguía-Rosas et al. 2011) that found that early flowering should be selected. It seems that the apparent high dependence of thrips for *C. pinnatifrons* floral resources is the main cause of the constant increase in the reproductive success that occurs throughout the flowering season. Further research should analyze the effect of such pattern on the progeny's fitness and the potential effect that antagonistic interactions could have over the individual's flowering phenology.

## ACKNOWLEDGMENTS

The authors would like to thank Asociación de Guías y Scouts de Costa Rica for their permission to conduct the research at Campo Escuela Iztarú at Cerros La Carpintera. All field assistants for the valuable help.

## SUPPORTING INFORMATION

Marquis (1988) synchrony index:

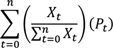

*X_t_* = number of flowering staminate or pistillate inflorescences at census *t*

*P_t_* = proportion of flowering staminate or pistillate individuals at census *t*

**Fig. S1.**
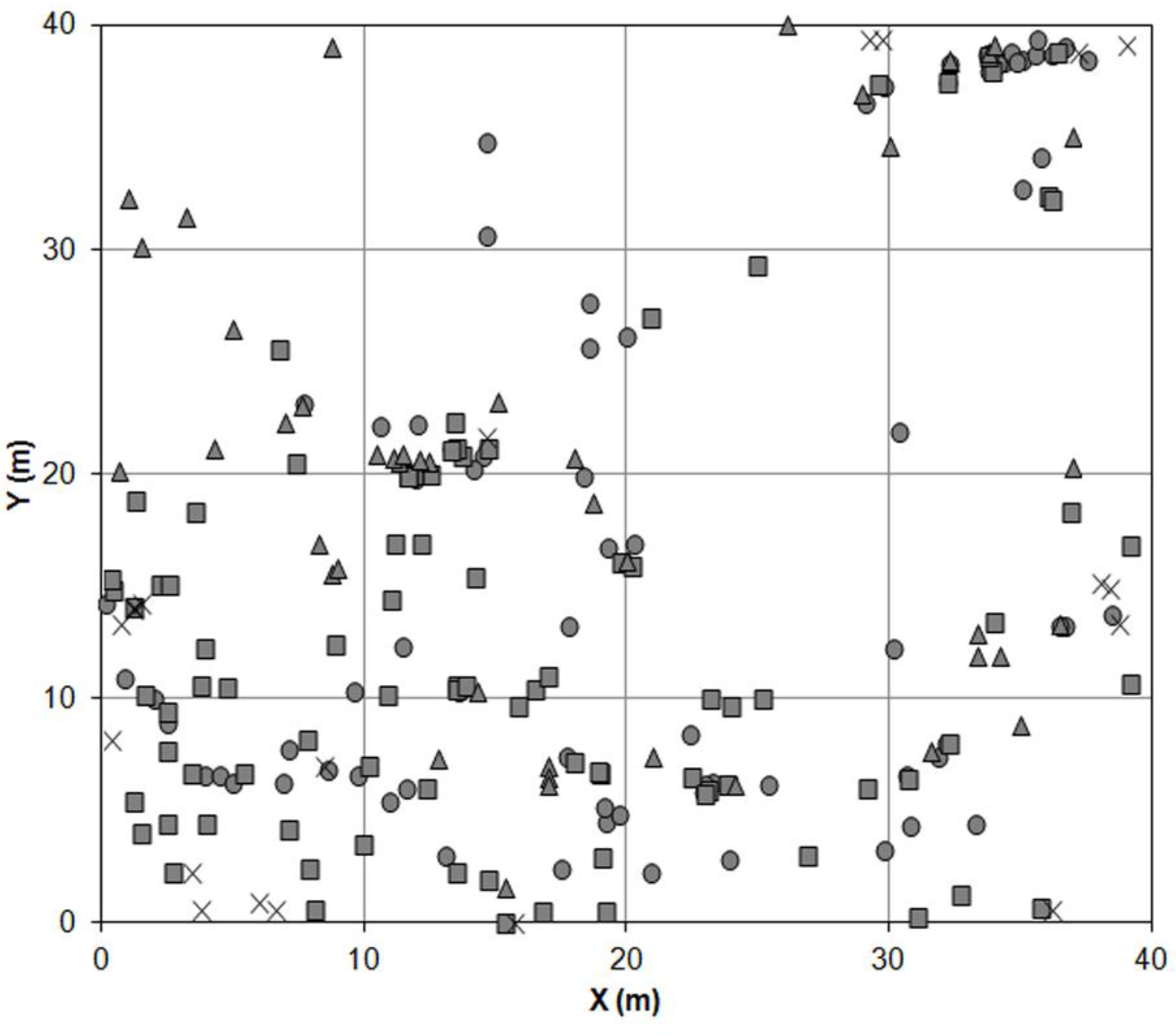
Study plot diagram. Circles and squares indicate pistillate and staminate individuals, respectively. Triangles indicate non reproductive individuals and crosses represent pistillate plants not used for reproductive success analyses.

**Fig. S2.**
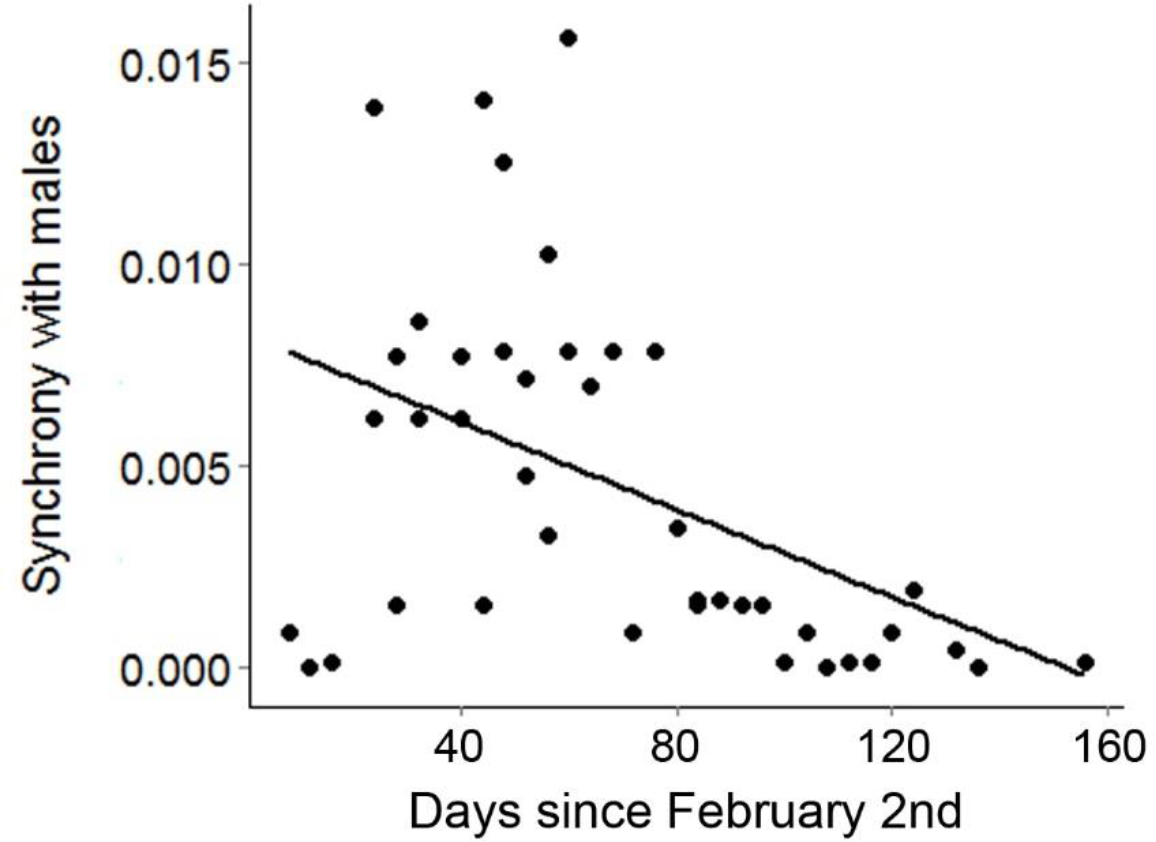
Interaction between census date and flowering synchrony with males during the 2012 season of *Chamaedorea pinnatifrons* (Arecaceae) at La Carpintera cloud forest, Costa Rica.

